# Lifespan regulation by targeting heme signaling in yeast

**DOI:** 10.1101/2024.01.20.576446

**Authors:** Praveen K. Patnaik, Nour Nady, Hanna Barlit, Ali Gülhan, Vyacheslav M. Labunskyy

## Abstract

Heme is an essential prosthetic group that serves as a co-factor and a signaling molecule. Heme levels decline with age, and its deficiency is associated with multiple hallmarks of aging, including anemia, mitochondrial dysfunction, and oxidative stress. Dysregulation of heme homeostasis has been also implicated in aging in model organisms suggesting that heme may play an evolutionarily conserved role in controlling lifespan. However, the underlying mechanisms and whether heme homeostasis can be targeted to promote healthy aging remain unclear. Here we used *Saccharomyces cerevisiae* as a model to investigate the role of heme in aging. For this, we have engineered a heme auxotrophic yeast strain expressing a plasma membrane-bound heme permease from *Caenorhabditis elegans* (ceHRG-4). This system can be used to control intracellular heme levels independently of the biosynthetic enzymes by manipulating heme concentration in the media. We observed that heme supplementation leads to significant lifespan extension in yeast. Our findings revealed that the effect of heme on lifespan is independent of the Hap4 transcription factor. Surprisingly, heme-supplemented cells had impaired growth on YPG medium, which requires mitochondrial respiration to be used, suggesting that these cells are respiratory deficient. Together, our results demonstrate that heme homeostasis is fundamentally important for aging biology and manipulating heme levels can be used as a promising therapeutic target for promoting longevity.

## Introduction

Heme is an essential iron-containing tetrapyrrole that serves both as a co-factor and a signaling molecule. As a protein prosthetic group, it is involved in different processes including electron transfer [1], chemical catalysis [2], and oxygen transport [3, 4]. It also serves as an important signaling molecule that regulates transcription factors controlling oxidative phosphorylation, antioxidant defense, and proliferation [5, 6]. Although heme is essential for health, it may also cause toxicity if present in excess [7]. Therefore, body heme levels are tightly regulated. Dysregulation of heme homeostasis has been implicated in the development of several diseases. In humans, defects in heme synthesis result in the accumulation of its intermediates leading to porphyria [8]. In addition, previous studies have shown that heme synthesis declines with age and its deficiency leads to mitochondrial dysfunction and iron accumulation leading to oxidative stress [9, 10]. On the other hand, deficiency of heme oxygenases, enzymes that are necessary for heme degradation, has been implicated in the development of several age-related disorders, such as Alzheimer’s disease, cancer as well as cardiovascular and metabolic diseases [11]. Moreover, recent genome-wide association studies (GWAS) have identified several genetic variants in genes involved in heme metabolism that are associated with human aging [12]. However, the molecular mechanisms by which heme regulates aging remain unclear.

Yeast *Saccharomyces cerevisiae* has proven to be a useful model for understanding the basic mechanisms by which dysregulation of heme homeostasis contributes to aging. Previous studies have shown that heme levels decrease with aging in yeast, which is associated with the decline of mitochondrial function [13]. The effects of heme on yeast lifespan were attributed to its direct functions as a co-factor in the electron transport chain in mitochondria and activating the heme activator protein (HAP) transcriptional complex which induces genes for maintaining mitochondrial biogenesis and function [14, 15]. Consistently, overexpression of Hap4, a component of the HAP complex, significantly extends lifespan [16]. Although increasing evidence suggests an important link between heme homeostasis and aging, the underlying mechanisms and causes of the decline in heme levels during aging are not completely understood.

In the present study, we developed a model to investigate the impact of heme on the yeast replicative lifespan. To distinguish the effects of endogenously produced heme from the effects of exogenous heme acquired from the media, we generated a heme auxotroph yeast strain, *hem1Δ*, which is unable to synthesize heme. Additionally, we introduced a membrane-bound heme permease from *C. elegans* into the yeast genome. This approach enabled us to directly control intracellular heme levels, independently of biosynthetic enzymes, by manipulating heme concentration in the media. Our findings demonstrate that supplementing heme, but not its precursor 5-aminolevulinic acid (ALA), is sufficient to extend lifespan in *S. cerevisiae*. Conversely, overexpression of the Hmx1 heme oxygenase, which is involved in heme degradation, results in a shortened replicative lifespan. Furthermore, we observed that heme supplementation extends lifespan independently of the Hap4 transcription factor, resulting in the inability of yeast cells to grow on respiratory carbon sources, indicating an adverse effect on mitochondrial function. Taken together, our findings reveal a crucial role of heme in the regulation of yeast replicative lifespan, laying the foundation for the development of heme-based therapeutic strategies for age-dependent diseases in human.

## Methods

### Yeast strains and culture conditions

The yeast strains used in this study and their genotypes are listed in Table S1. One-step polymerase chain reaction (PCR)-mediated gene disruption was performed using standard techniques to delete genes of interest. The genotypes of the resulting strains were verified using colony PCR.

To generate a yeast heme auxotroph strain expressing ceHRG-4, the sequence containing ceHRG-4 was first PCR amplified from pYES-DEST52-ceHRG-4 plasmid (pPP96) [17] and integrated into p416-GPD [18] vector using BamHI and XhoI restriction sites to generate pPP97 plasmid. The sequence encoding ceHRG-4 under the control of the glyceraldehyde-3-phosphate dehydrogenase promoter (*GPD1pr-ceHRG-4*) along with the URA3 marker were then amplified from pPP97 and integrated into the *hem1Δ* strain using homologous recombination at *URA3* locus. The resulting transformants were selected on SC-URA plates and verified by colony PCR and sanger sequencing.

To generate the *HMX1-OE* strain expressing *HMX1* under the control of the *ADH1* promoter (*ADH1pr-HMX1*), a 705 bp sequence of the *ADH1* promoter was amplified by PCR from yeast genomic DNA and integrated into the pRS306 plasmid using NotI and XhoI restriction sites. Subsequently, the fragment encompassing the URA3 marker and the *ADH1* promoter was amplified via PCR using the oPP399 and oPP400 oligonucleotides. These oligonucleotides have homology to the 3’ end of the *HMX1* promoter and the 5’ end of the HMX1 ORF, respectively. The resulting PCR product was then utilized for genomic integration. To create the yeast strain encoding an HA-tagged Hap4 protein, HA-tag sequence along with KanMX4 marker was amplified from pPP111 plasmid using the primers flanked on the 5’ side by sequence homologous to *HAP4* ORF and the 3’ side by sequence homologous to *HAP4* 3’-UTR for genomic integration. The resulting transformants were selected on YPD plates containing 200 µg/mL G418 and verified by colony PCR and sanger sequencing. The plasmids used in this study and sequences of primers used for generating yeast strains are listed in Table S2 and Table S3.

Yeast strains were cultured at 30°C in standard YPD medium (1.0% yeast extract, 2.0% peptone, and 2.0% glucose) unless otherwise stated. *hem1Δ* cells were maintained in YPD supplemented with 250 µM 5-aminolevulinic acid (ALA). To test the effect of exogenous heme on cell growth and replicative lifespan, the media was supplemented with hemin chloride. The stock hemin chloride solution was prepared by dissolving hemin in 0.3M NH_4_OH. The pH was adjusted to 8 with 6N HCl and the mixture was filter-sterilized through a 0.2 μm filter.

### Spot assays

The spot assays were used to assess the growth of strains under various growth conditions. Each strain to be tested was inoculated into a liquid culture medium and allowed to grow to the exponential phase. Once the cultures reached OD_600_ = 0.6, 10x serial dilutions for each strain were spotted on YPD agar plates (containing 2% glucose) or YPG plates (contaminating 3% glycerol) supplemented with indicated concentrations of ALA or hemin. The plates were incubated at 30°C for 48 hours before they were imaged.

### Replicative lifespan analysis

The cells were cultured on freshly prepared YPD plates at 30°C. Cells were monitored for cell divisions and the subsequent daughter cells were removed using a micromanipulator. The replicative lifespan was determined by counting the number of divisions each mother cell underwent before it ceased dividing. The lifespan assays were conducted in triplicates, and the data from three separate biological replicates were combined. The number of cells assayed, and statistical analysis of the lifespan data are shown in Table S4.

### RT-qPCR

Total RNA was extracted using hot acid phenol method followed by purification using Direct-zol RNA Miniprep Kit (Zymo Research). The RNA was treated with DNaseI, and 2 µg of RNA was used for cDNA synthesis using SuperScript III reverse transcriptase (Thermo Fisher Scientific) with random hexamer primers according to the manufacturer’s instructions. To analyze mRNA expression, real-time PCR was performed using SYBR Fast qPCR Master Mix (Kapa Biosystems) and the CFX-96 Touch Real-Time PCR Detection System (Bio-Rad Laboratories). To normalize the gene expression, *ACT1* was used as the reference gene. The primers used for RT-qPCR are listed in Table S3. Results are represented as means ± SEM from at least three independent experiments.

### Western blot analysis

A single colony was inoculated in 3 mL of YPD medium and incubated overnight at 30°C. The next day, the cells were diluted to an OD_600_ of 1 in 10 mL of fresh medium and grown until they reached the log phase. The cells were collected by centrifugation, washed with sterile water, and the pellet was frozen in liquid nitrogen and stored at -80°C for later use. For protein extraction, the cell pellet was re-suspended in 300 µL of Lysis Buffer (50 mM Tris pH 7.5, 150 mM NaCl, 1mM EDTA, 1% DMSO) containing 1mM phenylmethylsulfonyl fluoride (PMSF). The samples were homogenized with glass beads by vortexing at maximum speed for seven 30-second cycles and centrifuged for 5 minutes at 12,000 rpm at 4°C to obtain the supernatant. Equal amounts of proteins were resolved in 10% SDS-PAGE gels and transferred to PVDF membrane. The membrane was then probed with horseradish peroxidase (HRP)-conjugated HA tag monoclonal antibody (2-2.2.14) (ThermoFisher) using 1:1,000 dilution and signal quantification was performed using ImageJ software. Mouse anti-Pgk1 monoclonal antibody (1:5,000, Life Technologies) and HRP-conjugated secondary anti-mouse antibody (1:10,0000, Santa Cruz Biotechnology) were used to detect Pgk1 protein levels as a loading control.

### Heme quantification

Intracellular heme levels were assessed using the oxalic acid method as previously described with minor changes [19]. Briefly, overnight yeast cultures were diluted to OD_600_ = 0.2 units/mL and grown until cells reached OD_600_ = 0.8 units/mL. Subsequently, 8 OD_600_ units of cells were harvested by centrifugation at 2500 x g, washed with distilled water, and the pellet was resuspended in 500 µL of 20 mM oxalic acid. An additional 500 µl of 2 M oxalic acid was added, and the suspension was divided equally into two tubes. One set of sample tubes was placed in a heating block at 100° C for 45 min, while the other set of samples was kept at room temperature for the same duration as a baseline. Following incubation, samples were cooled to room temperature, and 100 µL of suspension was transferred per well into a black-well 96-well plate in duplicates. The fluorescence of porphyrin was measured using a Biotek plate reader with 400 nm excitation and 662 nm emission. Baseline values (from parallel unheated samples in oxalic acid) were subtracted, and the relative fluorescence intensity in arbitrary units (A.F.U.) was plotted.

### Quantification and Statistical analysis

Statistical analysis was performed using Prism 9.3.1 (GraphPad Software, Inc). Statistical significance of the heme quantification and RT-qPCR data was determined by calculating p values using one-way ANOVA. Error bars represent standard errors of the mean (SEM). The statistical significance of the lifespan data was evaluated using the Wilcoxon Rank-Sum test [20].

## Results

### Expression of *C. elegans* HRG-4 enhances heme import in yeast

The ability of yeast cells to synthesize heme might complicate the examination of how externally supplemented heme regulates the lifespan of *S. cerevisiae*. In order to avoid the effect of endogenously produced heme, we first introduced heme auxotrophy in yeast by deleting the *HEM1* gene encoding for the first enzyme of the heme biosynthesis pathway (**Figure 1A**). Consistent with prior reports, cells lacking *HEM1* exhibited impaired growth unless supplemented with 5-aminolevulinic acid (ALA) or an excess of exogenous heme (provided as hemin chloride) in the growth medium [21]. Because *S. cerevisiae* utilizes exogenous heme inefficiently, we introduced a gene encoding a plasma membrane-bound permease from *C. elegans* (ceHRG-4) into the yeast genome (**Figure 1B**). The growth of *hem1Λ1* cells expressing ceHRG-4 was rescued by lower concentration of heme (10 µM) compared to the *hem1Λ1* mutant, indicating a more efficient import of heme into the yeast cells in the presence of ceHRG-4 expression. Together, our data suggest that expressing ceHRG-4 increases heme uptake in yeast, allowing for precise control of intracellular heme levels by manipulating heme concentration in the media independently of heme biosynthetic enzymes.

**Figure 1.**
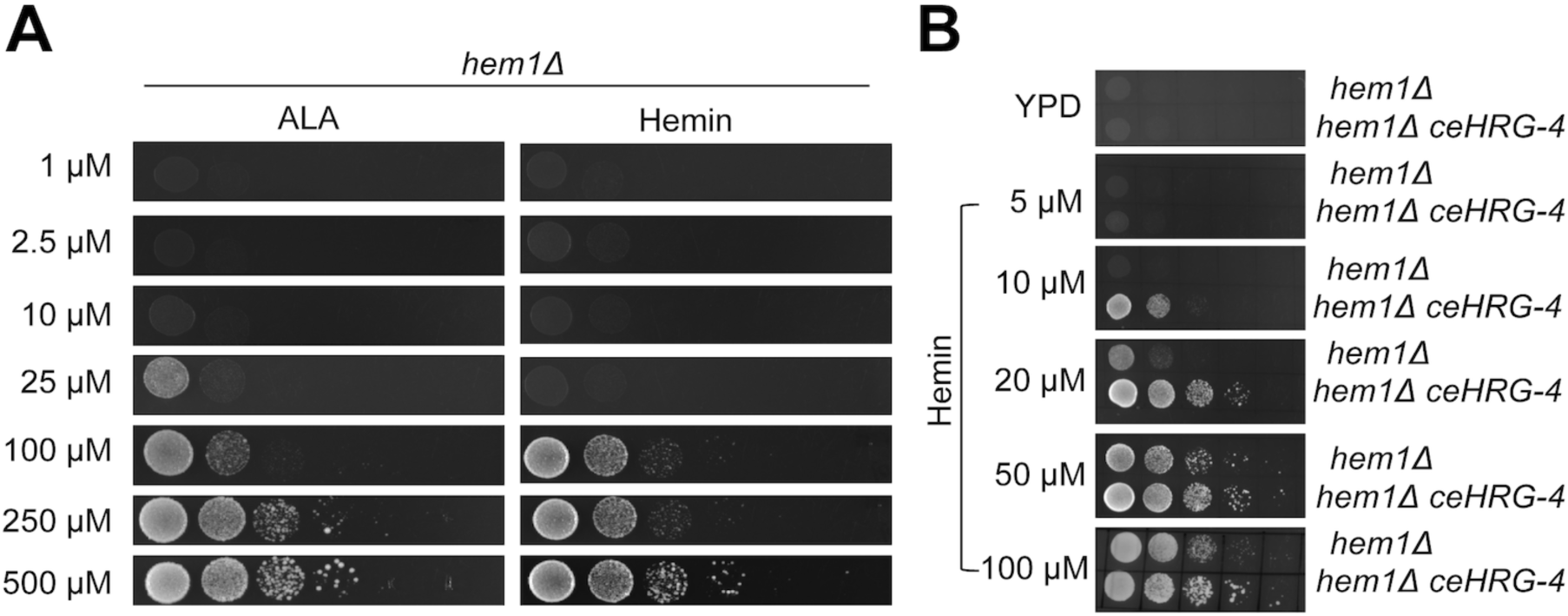
Engineering a heme auxotroph yeast strain to study the role of heme in aging. **A**) Cells lacking *HEM1* have impaired growth in the absence of ALA or hemin. The *hem1Δ* strain was cultured in YPD media containing 250 µM ALA overnight and serial (10x) dilutions were spotted on YPD plates supplemented with the indicated concentrations of ALA or hemin. Plates and were incubated at 30°C for 2 days prior to imaging. **B**) Expression of ceHRG-4 enhances heme import in yeast. The *hem1Δ* and *hem1Δ ceHRG-4* strains were cultured in YPD media containing 250 µM ALA overnight and serial (10x) dilutions were spotted on YPD plates supplemented with the indicated concentrations of hemin. A YPD plate without heme was used as a control. Plates were incubated at 30°C for 2 days prior to imaging.

### Heme supplementation extends yeast replicative lifespan

To investigate how heme levels affect aging, we analyzed yeast replicative lifespan in *hem1Λ1* cells expressing ceHRG-4 in the presence of different concentrations of heme in the media. Although *hem1Λ1*-*ceHRG-4* cells are unable to grow in the absence of heme, we found that increasing heme concentration from 10 to 100 µM extended lifespan by 67.8% (p<0.001) (**Figure 2A**). Similarly, wild-type cells expressing ceHRG-4 exhibited significant lifespan extension (46.3%; p<0.0001) when supplemented with 100 µM heme (**Fig 2B**). However, supplementing yeast with 5-aminolevulinic acid (ALA), a heme biosynthesis precursor, did not result in statistically significant difference in lifespan (**Figure 2C**). To validate that the effect of heme on lifespan is mediated by increased heme levels, we measured intracellular heme concentration in cells supplemented with different concentrations of hemin and ALA (**Figure 2D**). We found that supplementing heme to the media resulted in drastic increase in heme levels in cells expressing *ceHRG-4*. Conversely, ALA supplementation did not significantly alter intracellular heme levels, even at high concentrations.

**Figure 2.**
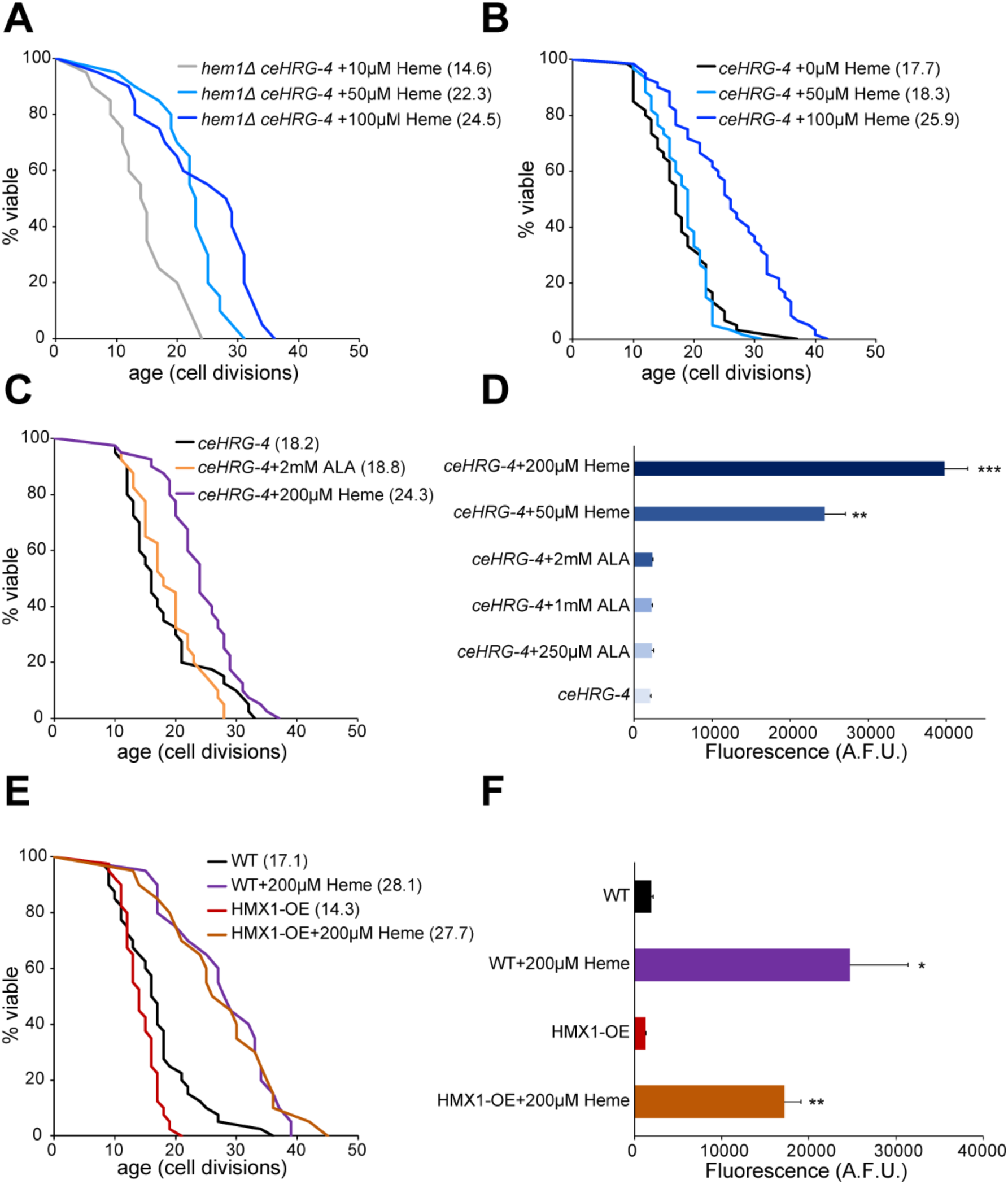
Heme supplementation extends yeast replicative lifespan. **A**) Supplementing heme extends the replicative lifespan in *hem1Δ* cells expressing ceHRG-4. Lifespan of the *hem1Δ ceHRG-4* mutant was analyzed on YPD plates supplemented with indicated concentrations of hemin. The average lifespan is shown in parentheses. **B**) Replicative lifespan of wild-type cells expressing ceHRG-4. **C)** Supplementing 5-aminolevulinic acid (ALA) does not extend replicative lifespan in yeast. **D**) Spectrofluorometric quantification of intracellular heme levels in wild-type cells expressing *ceHRG-4* supplemented with indicated concentrations of hemin and ALA. Error bars represent SEM of three biological replicates, each containing two technical replicates. **p<0.01, ***p<0.001 compared with *ceHRG-4* control (one-way ANNOVA). **E**) Shortened lifespan in cells overexpressing HMX1 (HMX1-OE) can be rescued by heme supplementation. **F**) Increased lifespan in yeast cells supplemented with heme correlates with intracellular heme concentration. Error bars represent SEM of three biological replicates, each containing two technical replicates. *p<0.05, **p<0.01 compared with wild-type control (one-way ANNOVA).

Given the beneficial effect of heme supplementation on lifespan, we sought to investigate the effects of decreasing intracellular heme levels on lifespan. To this end, we overexpressed the *HMX1* gene, which encodes heme oxygenase involved in intracellular heme degradation. *HMX1* overexpression (*HMX1-OE*) decreased lifespan 16.4% compared to wild-type cells (**Figure 2E**). Furthermore, supplementing the *HMX1-OE* strain with heme was able to rescue the shortened lifespan caused by the *HMX1* overexpression, and this increase in lifespan correlated with intracellular heme concentration (**Figure 2F**). Together, these findings suggest that heme plays a critical role in the regulation of replicative lifespan in yeast.

### Heme extends yeast lifespan independently of Hap4

Previous studies have shown that supplementation of yeast cells with heme or its precursors results in an increased expression of the Hap4 transcription factor [22]. Hap4 plays a key role in the regulation of mitochondrial function and respiration in yeast [23, 24]. Furthermore, overexpression of *HAP4* has been shown to extend the yeast lifespan [16]. Based on these observations, we hypothesized that the extended lifespan resulting from heme supplementation could be attributed to the overexpression of Hap4. To test this hypothesis, we first asked whether heme supplementation would lead to increased Hap4 expression. For this, we quantified the levels of the Hap4 protein in the cells treated with different concentration of heme using Western blot analysis (**Figure 3A**). Contrary to expectations, we found that Hap4 protein levels were significantly decreased (p<0.001) in the presence of heme supplementation compared to the control cells.

**Figure 3.**
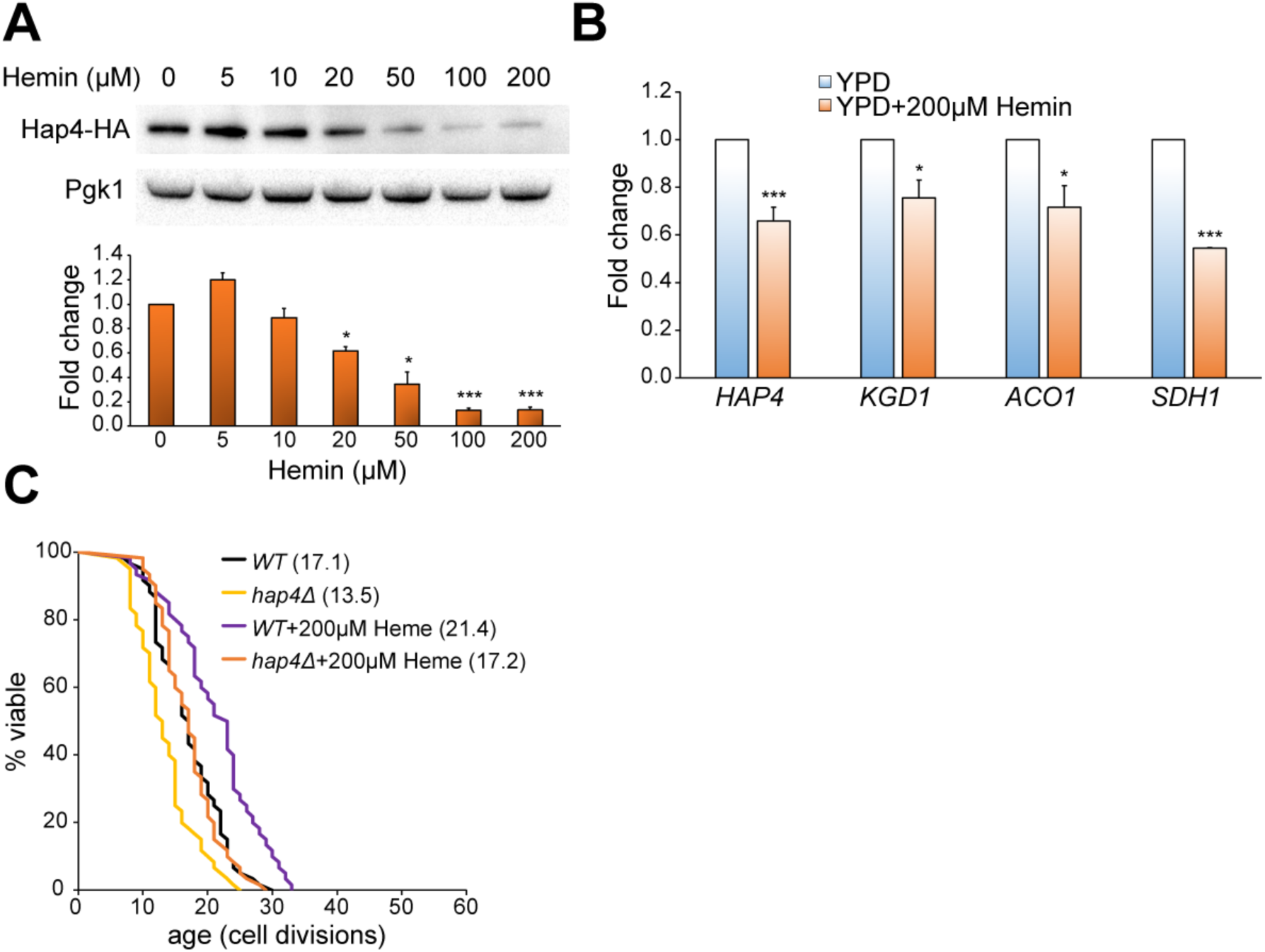
Heme supplementation inhibits expression of Hap4. **A**) Hap4 protein levels are repressed by heme. The expression of Hap4-HA in cells treated with different concentrations of heme was assessed by Western blotting using an anti-HA antibody. Pgk1 was used as a loading control. Quantification of protein expression was carried out using ImageJ software. Error bars represent SEM of three independent experiments. **B**) The expression of *HAP4* and its targets in exponentially growing cells was assessed using RT-qPCR. Error bars represent SEM of three independent experiments. * p<0.05; *** p<0.001 (one-way ANOVA). **C**) Heme supplementation extends replicative lifespan independently of *HAP4*. Lifespan of the wild-type (*WT*) and *hap4Δ* cells was analyzed on YPD plates in the presence or absence of 200 µM hemin. The average lifespan is shown in parentheses.

It has been previously shown that Hap4 is susceptible to proteasome-mediated degradation [25]. To test whether decreased Hap4 protein levels can be attributed to its increased proteasomal degradation, we treated cells with 200 µM heme in the presence or absence of 75 µM MG132 proteasome inhibitor [26]. Because *S. cerevisiae* is resistant to MG132 [27], we employed the *erg6Δ* strain with increased permeability to proteasome inhibitors [28]. When *erg6Δ* cells were supplemented with 200 µM heme, we observed a significant decrease in Hap4 protein (**Supplementary Figure S1**). However, treatment with MG132 was not able to restore protein levels, suggesting that the decrease in Hap4 levels is not due to proteasomal degradation. Furthermore, we analyzed abundance of *HAP4* mRNA and expression of the Hap4 transcription factor targets, including *KGD1*, *ACO1*, and *SDH1* (**Figure 3B**) using RT-qPCR. We observed a significant decrease in *HAP4* mRNA levels (p<0.001) and diminished expression of the Hap4 targets when yeast cells were exposed to a treatment with high heme concentration.

To directly test if the increased lifespan in response to heme supplementation requires Hap4, we analyzed replicative lifespan in wild-type cells and the *HAP4* deletion mutant (*hap4Δ*) in the absence and presence of heme supplementation (**Figure 3C**). Although the deletion of *HAP4* resulted in a shortened replicative lifespan (p<0.001) compared to the wild-type strain, the supplementation of *hap4Δ* with heme was sufficient to significantly extend the lifespan. Taken together, our findings suggest that heme supplementation causes a reduction in *HAP4* expression, and that the lifespan extension effect of heme cannot be attributed to the increased activity of the Hap4 transcription factor.

### Heme supplementation negatively affects respiration

Since Hap4 is involved in regulating mitochondrial biogenesis in yeast [23, 24], we investigated whether the heme-induced reduction in *HAP4* expression would result in decreased cell respiration. To explore this, we examined the growth of yeast cells supplemented with varying heme concentrations on YPG plates containing glycerol as a carbon source that requires respiration to be utilized (**Figure 4A**). Notably, we observed a decline in yeast growth on YPG plates as heme concentration increased. Additionally, heme supplementation inhibited cellular growth when cells were cultured in media containing alternative carbon sources that rely on respiration, including ethanol and lactate (**Figure 4B**). We further examined the impact of heme supplementation on population doubling time in glucose-containing medium (YPD). Our data revealed that ceHRG-4 expressing cells treated with heme exhibited normal growth during exponential phase (fermentative growth) on YPD medium containing 2 % glucose (**Figure 4C**). However, after glucose was exhausted during diauxic shift, these cells were unable to switch metabolism from fermentation to respiration and utilize the produced ethanol as a carbon source, leading to impaired growth compared to untreated cells. Together, our findings indicate that heme supplementation negatively affects Hap4 expression and results in decreased respiratory capacity.

**Figure 4.**
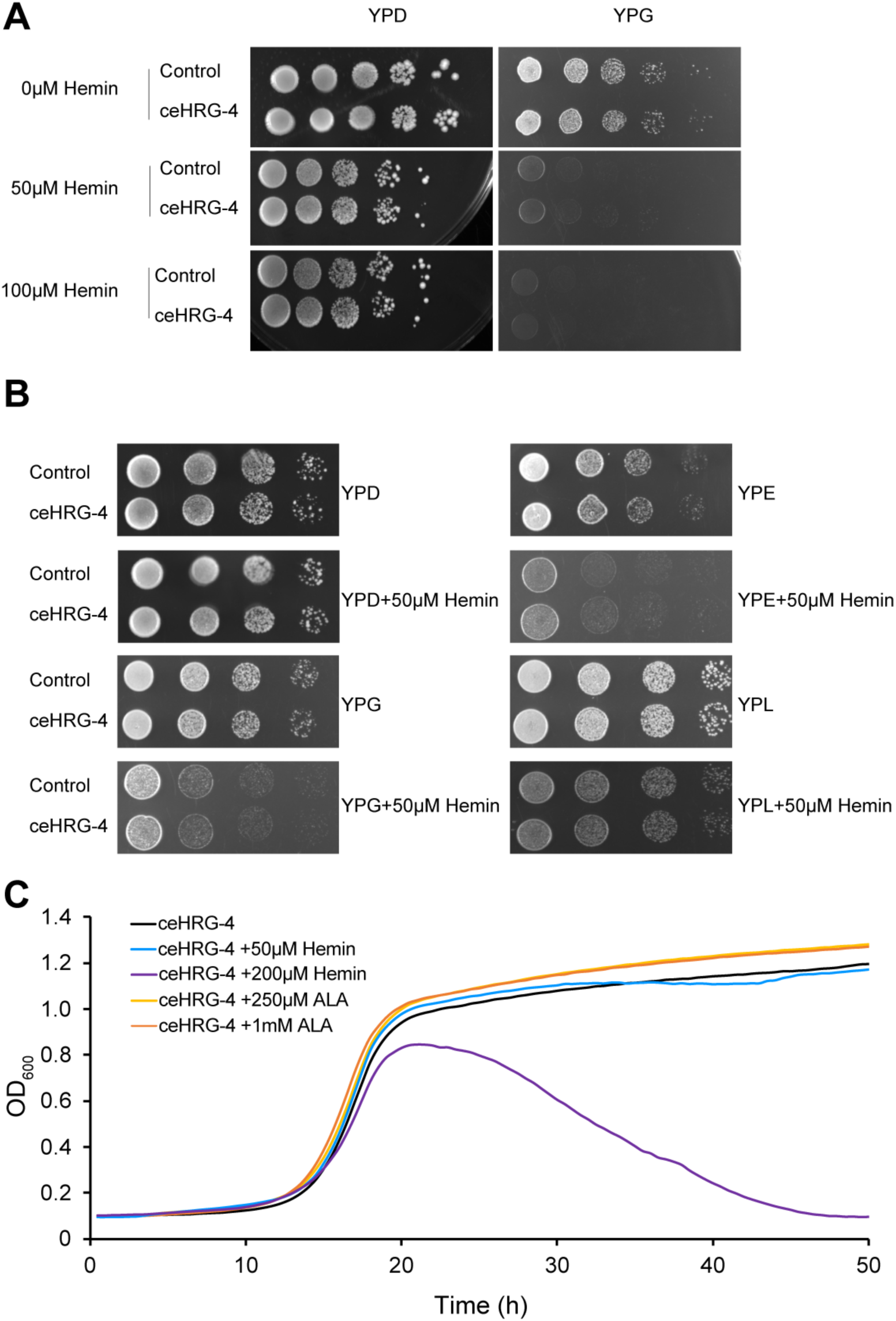
Heme supplementation negatively affects respiration. **A**) Supplementation of hemin leads to delayed growth of wild-type and ceHRG-4 expressing cells on glycerol-containing medium (YPG). Yeast strains were cultured in YPD media overnight and serial (10x) dilutions were spotted on YPD and YPG plates containing indicated concentrations of hemin. Plates were incubated at 30°C for 2 days prior to imaging. **B**) Heme supplementation inhibits cellular growth in media containing alternative carbon sources that rely on respiration. Yeast strains were cultured in YPD media overnight and serial (10x) dilutions were spotted on agar plates with YPD (2% glucose), YPG (3% glycerol), YPE (5% ethanol), and YPL (3% lactate) media in the presence or absence of 50 µM hemin. Plates were incubated at 30°C for 2 days prior to imaging. **C**) Heme supplementation leads to the growth inhibition of *ceHRG-4* expressing cells during respiratory phase in YPD media.

## Discussion

Heme is an essential prosthetic group for enzymes involved in multiple biological functions. Defects in heme synthesis and transport have been associated with multiple human disorders, including anemia and mitochondrial dysfunction. However, the causes of the decline in heme levels during aging and mechanisms by which heme regulates lifespan are not understood. In this study, we used the *S. cerevisiae* model to investigate the role of heme in aging. To tightly regulate heme concentration, we induced heme auxotrophy in yeast through the deletion of the *HEM1* gene. Since budding yeast utilizes exogenous heme inefficiently even in the absence of endogenous heme synthesis [21, 29], we employed a previously established strategy to manipulate heme uptake in yeast by overexpressing a plasma membrane-bound heme permease from *C. elegans* (ceHRG-4) [30].

How do intracellular heme levels affect lifespan in yeast? To distinguish the effects of endogenously produced heme from the effects of exogenous heme, we supplemented yeast cells with either ALA, a precursor of heme biosynthesis, or directly provided hemin in the media. Our findings revealed that hemin supplementation significantly extends the yeast replicative lifespan. In contrast, supplementing ALA, which requires mitochondria to be utilized for the synthesis of intracellular heme, did not significantly affect lifespan. The observation that ALA supplementation did not extend lifespan may be attributed to low intracellular heme levels, possibly resulting from inefficient ALA transport or negative feedback regulation of heme biosynthesis enzymes.

Among the potential causes of the decline in heme levels during aging are increased heme degradation and dysregulation of heme biosynthesis. Using yeast as a model, we have recently shown that expression of *HMX1*, involved in heme degradation, is increased with aging whereas deletion of the *HMX1* gene leads to lifespan extension [31]. The yeast *HMX1* gene is regulated by the Aft1 transcription factor. During aging, increased activity of the Aft1 leads to increased expression of *HMX1* resulting in decreased intracellular heme levels. Consistent with the role of Hmx1 in heme degradation, our data show that overexpression of *HMX1* shortens lifespan (**Figure 2**).

Furthermore, the rate of heme production declines with aging. For example, the activity of the ALA synthase [32], a rate-limiting enzyme in heme biosynthesis, and ALA levels have been shown to decrease with aging [33, 34]. Notably, ALA supplementation was able to restore muscle function and extend healthspan in *Drosophila melanogaster* [35]. Additionally, the mRNA-binding protein Cth2, which is involved in the regulation of heme biosynthesis enzymes, is aberrantly expressed with aging leading to the repression of heme synthesis [31]. On the other hand, mitochondria play a key role in heme biosynthesis. It is possible that reduced heme biosynthesis during aging is a consequence of mitochondrial disfunction in aging cells.

The effects of heme on yeast lifespan can be attributed to its direct function as a cofactor in the electron transport chain (ETC) in mitochondria or serving as a signaling molecule. We hypothesized that heme could extend lifespan by activating the Hap4 transcription factor, which induces expression of genes required for maintaining mitochondrial biogenesis and function. Surprisingly, we found that Hap4 levels were decreased in cells treated with hemin. The decrease in Hap4 protein levels was not due to proteasomal degradation, but instead was driven by changes in *HAP4* mRNA abundance leading to inhibition of Hap4 activity and inability of cells to grow on respiratory carbon sources. We also found that heme supplementation extends lifespan independently of the Hap4 transcription factor. Since Hap4 expression is downregulated by heme and *HAP4* is dispensable for its lifespan extension effects, these results suggest that heme supplementation may extend lifespan through alternative mechanisms, such as increased activity of enzymes requiring heme as a cofactor or activation of adaptive stress responses.

Previous studies in model organisms have shown that inhibition of mitochondrial function as well as components of the ETC and TCA cycle can extend lifespan in several species [36], including yeast [37], worms [38], flies [39], and mice [40]. For example, downregulation of aconitase and isocitrate dehydrogenase, two enzymes in the TCA cycle, leads to lifespan extension in worms [41]. Additionally, a genetic screen in yeast measuring replicative lifespan of 4698 deletion mutants, identified mitochondrial translation and components of TCA among the most enriched functional groups [42]. However, specific genes and stress response pathways that connect these changes to longevity remain unknown. Further studies analyzing age-dependent changes in transcriptional network of cells in response to heme treatment may provide additional insights into pathways associated with lifespan extension.

Together, our results suggest the existence of a previously unanticipated role of heme in the regulation of lifespan. Our findings raise an exciting possibility that manipulating heme levels by genetic or pharmacological means can be used as a therapeutic strategy for age-related diseases in humans.

## Acknowledgments

We would like to thank Dr. Iqbal Hamza for providing the pYES-DEST52-ceHRG-4 plasmid used in this study. This work was supported by NIH Grants AG058713 and AG066704 to VML.

## Competing interests

The authors declare no competing interests.

## Supplementary Materials

**Supplementary Figure S1:** Heme supplementation does not affect the proteasomal degradation of Hap4

**Supplementary Table S1:** Yeast strains used in this study

**Supplementary Table S2:** Plasmids used in this study

**Supplementary Table S3:** Oligonucleotides used in this study

**Supplementary Table S4:** Statistical analysis of lifespan experiments

## Notes

### Competing Interest Statement

The authors have declared no competing interest.

