## Supplementary Figure S1 for "Lifespan regulation by targeting heme signaling in yeast"

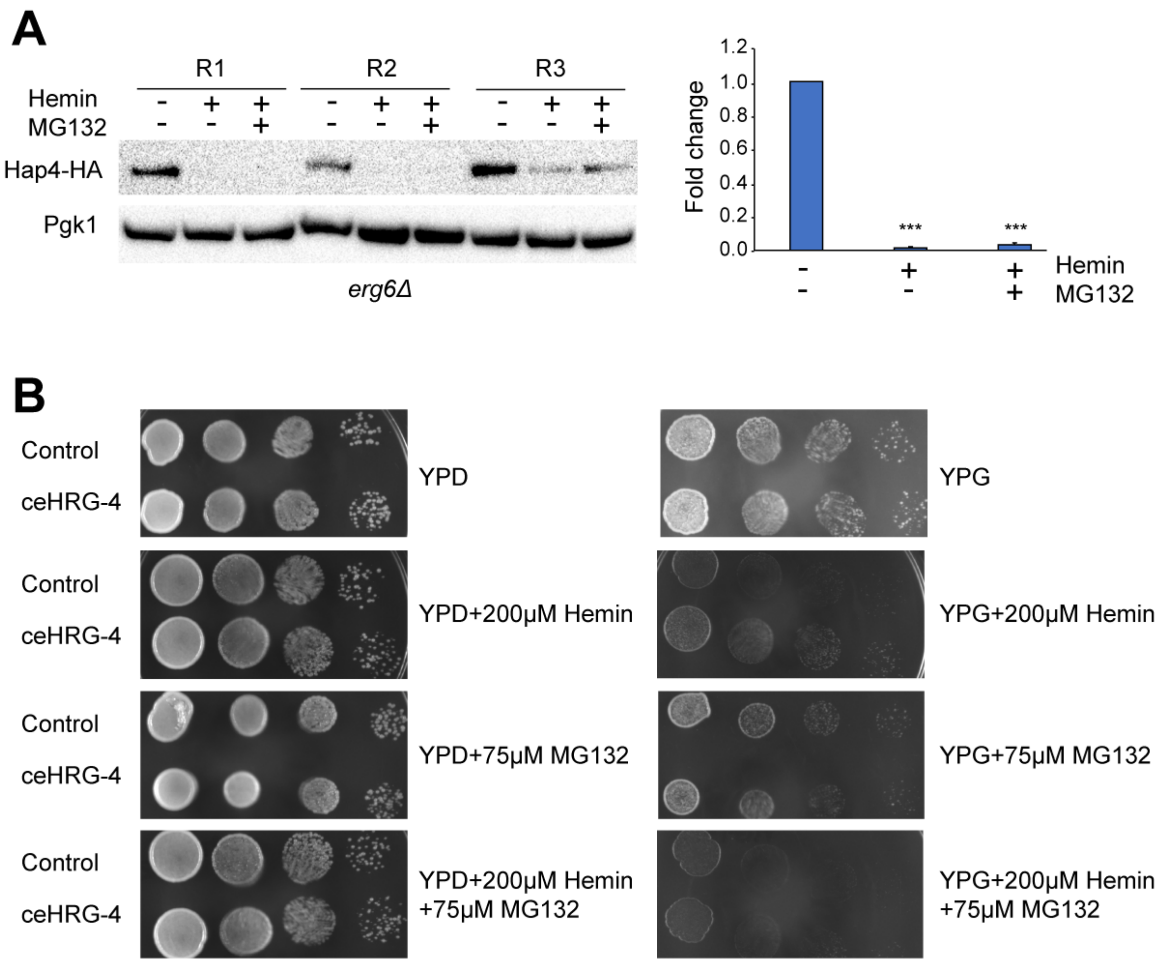


**Supplementary Figure S1. Heme supplementation does not affect the proteasomal degradation of Hap4. A**) Western blot quantification of Hap4-HA protein in *erg6Δ* strain. Yeast cells were treated with either 200 $\mu$M heme, 75 $\mu$M MG132 (proteasome inhibitor) or both. Pgk1 was used as a loading control. Right panel shows quantification of protein expression using ImageJ software. Error bars represent SEM of three independent experiments. ***, $p<0.001$ (one-way ANOVA). **B**) The proteasomal inhibitor does not rescue the growth of wild-type and ceHRG-4 expressing *erg6Δ* cells treated with heme on glycerol-containing medium (YPG). Yeast strains were cultured in YPD media overnight and serial (10x) dilutions were spotted on YPD and YPG plates containing 200 $\mu$M hemin, 75 $\mu$M MG132, or both. Plates were incubated at 30°C for 2 days prior to imaging.
