## Supplementary Table S1 for "Lifespan regulation by targeting heme signaling in yeast"

**Supplementary Table S1. Yeast strains used in this study.**

| Identifier | Genotype | Source |
|---|---|---|
| PP109 (BY4741) | *MATa his3Δ1 leuΔ0 met15Δ0 ura3Δ0* | Horizon Discovery |
| PP408 | *MATa his3Δ1 leu2Δ0 met15Δ0 ura3Δ0 HMX1::ADH1pr-HMX1-URA3* | This study |
| PP438 | *MATa his3Δ1 leuΔ0 met15Δ0 ura3Δ0 hem1Δ::KanMX4* | This study |
| PP454 | *MATa his3Δ1 leuΔ0 met15Δ0 hem1Δ::KanMX4 ura3Δ::pGAP-ceHRG-4-HA-URA3* | This study |
| PP473 | *MATa his3Δ1 leuΔ0 met15Δ0 ura3Δ::GPDpr-ceHRG-4-HA-URA3* | This study |
| PP502 | *MATa his3Δ1 leuΔ0 met15Δ0 ura3Δ::GPDpr-ceHRG-4-HA-URA3 erg6Δ::NatMX4* | This study |
| PP505 | *MATa his3Δ1 leuΔ0 met15Δ0 ura3Δ0 hap1Δ::KanMX4* | This study |
| PP507 | *MATa his3Δ1 leuΔ0 met15Δ0 ura3Δ0 hap4Δ::HAP4-HA-KanMX4* | This study |
