## Supplementary Table S2 for "Lifespan regulation by targeting heme signaling in yeast"

**Supplementary Table S2. Plasmids used in this study.**

| Plasmid | Description | Source |
|---|---|---|
| pPP111 | pDZ415-HA | This study |
| pPP52 | pRS306-ADH1pr | This study |
| pPP97 | p416-GPD-HRG4 | This study |
