## Supplementary Table S3 for "Lifespan regulation by targeting heme signaling in yeast"

**Supplementary Table S3. Oligonucleotides used in this study.**

| Oligonucleotides | Sequence | Purpose |
|---|---|---|
| oPP085_ACT1_F | TCGTTCCAATTTACGCTGGTT | RT-qPCR |
| oPP086_ACT1_R | CGGCCAAATCGATTCTCAA | RT-qPCR |
| oPP228_ACO1_F | GACCATTTTCACTGTTACTCC | RT-qPCR |
| oPP229_ACO1_R | GATATCTCTACGATCCCATTG | RT-qPCR |
| oPP238_KGD1_F | GATAAGAGGTTCGGTTTAGAAG | RT-qPCR |
| oPP239_KGD1_R | GTTTACGGACCACATTGGAT | RT-qPCR |
| oPP260_HAP4_F | CTGATTCTCCAGCAGATTTC | RT-qPCR |
| oPP261_HAP4_R | CGTTATTCGTGTTGACTTTG | RT-qPCR |
| oPP268_Hap4pr_F | TCTCCTAGTACATCAAAGAGC | *HAP4* deletion |
| oPP269_Hap4orf_R | TAAAATGGTTACTACGAGGGC | *HAP4* deletion |
| oPP291_SDH1_F | AGAGGTGTTGGTAAGAAAAAG | RT-qPCR |
| oPP292_SDH1_R | GGGAATACCACCCATGTTAT | RT-qPCR |
| oPP378_TEFpr_R | CTGCAGCGAGGAGCCGTAAT | ceHRG-4 sequencing |
| oPP399_HMX1pr_Ura3pr_F | ACAGCATATATACACACACACACATAAAATAACCGCAAAAttcaattcatcatttttttt | *HMX1* overexpression |
| oPP400_HMX1orf_ADHpr_R | CAGTGGGTGAGGGTATGATTGTATTGCTACTGTCCTCCATagttgattgtatgcttggta | *HMX1* overexpression |
| oPP431_URA3_GAPpr_F | AAGGATAAGTTTTGACCATCAAAGAAGGTTAATGTGGCTGtcattatcaatactcgccat | ceHRG-4 integration |
| oPP432_URA3_p416_R | TTGGATAGTTCCTTTTTATAAAGGCCATGAAGCTTTTTCTtactgagagtgcaccatacc | ceHRG-4 integration |
| oPP437_Hap4_HA_F | CCTTGACGAAGATGTCGATTTTTTAAAGGTACAAGTATTTagcttaggtggaatgtaccc | Hap4-HA integration |
| oPP438_Hap4_HA_R | TTTTTAGTTGTTTTCGTTTTATTGCAACATGCCTATTTCAgcataggccactagtggatc | Hap4-HA integration |
| oPP458_ERG6_F | GATGCCGAAGAACGTCGTCT | *ERG6* genotyping |
| oPP459_ERG6_R | TACCACCCGGTTTCAAAACC | *ERG6* genotyping |
